# Hippocampal motifs

**DOI:** 10.1101/001636

**Authors:** Zahra M. Aghajan, Lavanya Acharya, Jesse Cushman, Cliff Vuong, Jason Moore, Mayank R. Mehta

**Affiliations:** W. M. Keck Center for Neurophysics, Integrative Center for Learning and Memory, and Brain Research Institute; Department of Physics and Astronomy, University of California at Los Angeles; Biomedical Engineering Interdepartmental Program, University of California at Los Angeles; Neuroscience Interdepartmental Program, University of California at Los Angeles; Departments of Neurology and Neurobiology, University of California at Los Angeles

## Abstract

Dorsal Hippocampal neurons provide an allocentric map of space^1^, characterized by three key properties. First, their firing is spatially selective^1–3^, termed a rate code. Second, as animals traverse through place fields, neurons sustain elevated firing rates for long periods, however this has received little attention. Third the theta-phase of spikes within this sustained activity varies with animal’s location, termed phase-precession or a temporal code^4–10^. The precise relationship between these properties and the mechanisms governing them are not understood, although distal visual cues (DVC) are thought to be sufficient to reliably elicit them^2^^,^^3^. Hence, we measured rat CA1 neurons’ activity during random foraging in two-dimensional VR—where only DVC provide consistent allocentric location information— and compared it with their activity in real world (RW). Surprisingly, we found little spatial selectivity in VR. This is in sharp contrast to robust spatial selectivity commonly seen in one-dimensional RW and VR^7–11^, or two-dimensional RW^1–3^. Despite this, neurons in VR generated approximately two-second long phase precessing spike sequences, termed “hippocampal motifs”. Motifs, and “Motif-fields”, an aggregation of all motifs of a neuron, had qualitatively similar properties including theta-scale temporal coding in RW and VR, but the motifs were far less spatially localized in VR. These results suggest that intrinsic, network mechanisms generate temporally coded hippocampal motifs, which can be dissociated from their spatial selectivity. Further, DVC alone are insufficient to localize motifs spatially to generate a robust rate code.

---

When an animal explores a two-dimensional environment, hippocampal neurons fire in a spatially selective fashion to form an allocentric map of space^1^. The mechanisms governing this selectivity remain to be understood. DVC are thought to be the primary factor governing hippocampal spatial selectivity^2^^,^^3^, although the contribution of other modalities have not been ruled out^12^^,^^13^. While these uncontrolled cues are difficult to eliminate in RW, they can be either removed or made spatially non-informative in VR. Hence, rodent hippocampal activity has been recently measured on one-dimensional mazes where neurons show comparable spatial selectivity in RW and VR^7–9,11^. Spatial selectivity on one-dimensional VR tracks could arise not only from DVC^2^^,^^3^, but also from self-motion cues^9^^,^^14^^,^^15^ or intrinsic network mechanisms^9,16–19^ because these are highly correlated with spatial location. To unequivocally determine the contribution of only DVC it is necessary to measure hippocampal activity in a two-dimensional VR where rats explore an environment randomly, as commonly done in RW^1–3^. This ensures that unlike on stereotypical trajectories, self-motion or internal cues do not have a fixed relationship with DVC and hence they do not provide reliable spatial information.

We measured hippocampal activity during a random foraging task in RW and VR^20^^,^^21^. In VR rats were body fixed with a harness on a floating ball, allowing head movements (see Methods)^9^^,^^21^. The two worlds had identical dimensions and DVC (Fig. 1a). In RW, visual and somatosensory cues indicated the platform edge. In VR, steps beyond the virtual edge of the platform caused no change in the visual scene. Rats quickly learned to avoid or turn away from the virtual edges based entirely on visual cues^21^. The amount of time they spent away from the edges and in the center of the platform was comparable in the two worlds, although they ran at somewhat lower speeds in VR (Fig. 1a). We recorded the activity of 585 (501) neurons in dorsal CA1 from three rats that were active in RW (VR) at mean firing rates above 0.2Hz during locomotion at speeds >5cm/s. Only these cells were used for all subsequent analysis unless otherwise stated. Neurons fired vigorously in restricted regions of space in RW as expected (Fig. 1b, c, Extended Data Fig. 1)^1^. In contrast they showed little spatial selectivity in VR (Fig. 1b, d, Extended Data Fig. 1).

**Figure 1.**
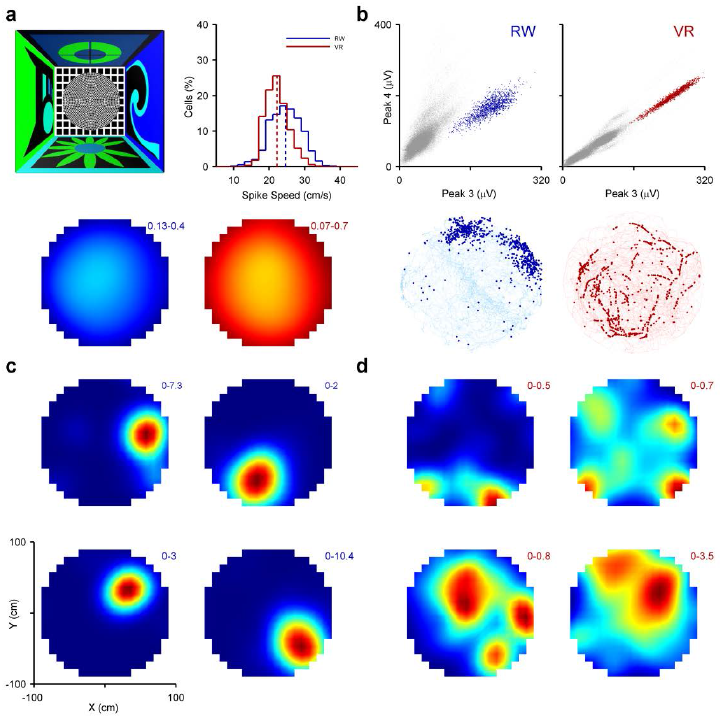
Similar rat behavior but different neural ratemaps in two-dimensional RW and VR. **a)** Top Left: Top-down view of the VR and RW mazes showing a 200cm diameter elevated platform centered in a 300×300cm room with distinct visual cues on the walls. Top right: Mean running speed at the time of occurrence of spikes (excluding speeds <5cm/s) was slightly reduced (8%, p<10^−10^) in VR (22.26±3.93cm/s, Red) compared to RW (24.12±4.81cm/s, Blue). Bottom: Percentage of time spent in all parts of the maze, averaged across all rats showing that rats spent comparable time away from edges in RW (left, blue) and VR (right, red). Numbers indicate range; lighter shades indicate higher values. These color conventions (RW, blue shades; VR, red shades; lighter shades, higher values) apply to all subsequent plots. **b)** Top: Scatterplots of spike amplitudes (grey dots) on one tetrode channel versus another in RW and VR. Colored dots are spikes from isolated neurons. Bottom: Position of the rat in RW and VR at the time of occurrence of spikes (darker dots) from the corresponding neurons (top) overlaid on the trajectory of the rat (lighter trace). **c)** Spatial ratemaps of four neurons in RW. Numbers indicate range (higher rates, hotter colors). **d)** Same as in panel C but in VR.

Across the ensemble, VR neurons had moderately reduced (29%) mean firing rates but the peak firing rates were greatly reduced in VR (64%) (Fig. 2a, Extended Data Fig. 2). Spatial information content also showed a large 79% reduction in VR (Fig. 2b, Extended Data Fig. 2). Spatial stability over the period of a session (see Methods) of VR ratemaps was also greatly reduced (90%) compared to stable and spatially localized RW ratemaps (Fig. 2c); VR ratemap stability was near chance level (Fig. 2c, d, Extended Data Fig. 3). Information content was negatively correlated with mean firing rate in both worlds (Fig. 2e), but cells with similar mean rates had much smaller spatial information content and stability in VR compared to RW. Also, stability increased with mean rate in RW but not VR (Extended Data Fig. 2). Thus the loss of spatial information and stability in VR was not due solely to the reduced mean firing rates.

**Figure 2.**
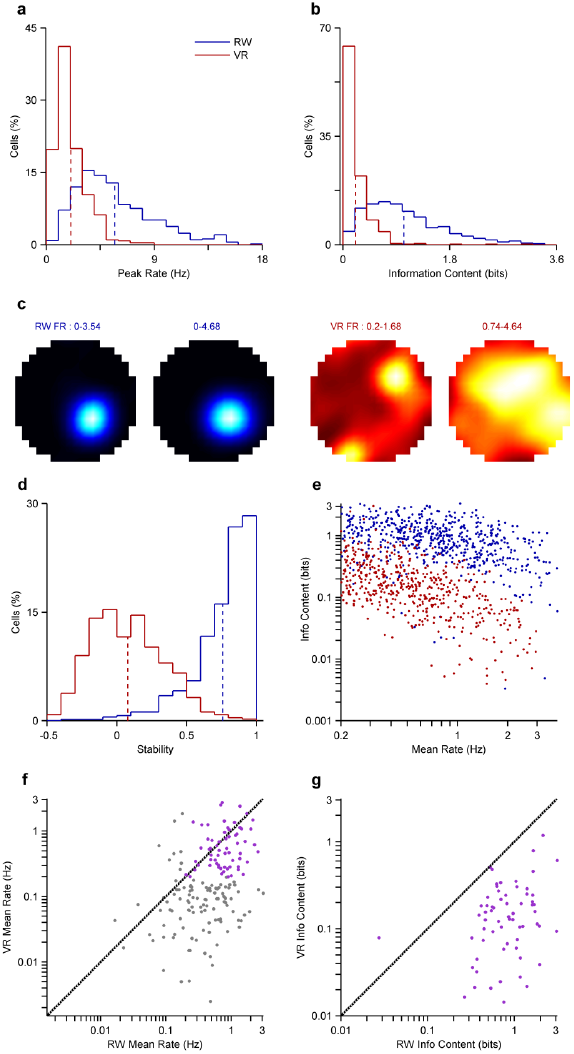
Reduced activity, spatial selectivity and stability of ratemaps in VR. **a)** Peak firing rates of neurons were 64% (p<10 ^−10^) smaller in VR (2.04±0.06Hz) compared to RW (5.72±0.13Hz). **b)** Spatial information content in VR (0.21±0.01bits) was 79% (p<10^−10^) lower than in RW (1.03±0.03bits). **c)** Ratemaps of a neuron during first and second halves of a session in RW and VR. **d)** Stability of ratemaps in VR (0.08±0.01) was 90% reduced (p<10^−10^) compared to RW (0.76±0.01). **e)** Spatial information was negatively correlated with the mean firing rate of a cell in both worlds (RW r = -0.41, p<10^−10^; VR r = -0.35, p<10^−10^) **f)** For cells recorded in both worlds on the same day mean firing rate was correlated regardless of minimum firing rate (r=0.31, p<10^−4^). This was also true for cells active in both environments (purple, r=0.36, p<0.05). **g)** Spatial information for cells active in both environments was also significantly correlated (r=0.38, p<10^−3^).

Was there any relationship between the activity of the same neuron in RW and VR? To address this we characterized the activity of 174 neurons recorded in RW and VR on the same day (Fig. 1b). For these neurons, the mean firing rates were significantly correlated between RW and VR (Fig. 2f). Of these, only 37% had a mean firing rate above 0.2Hz, referred to as “active,” in both worlds and were used for subsequent same-cell analyses. The spatial information in VR, although much lower, was significantly correlated to that in RW (Fig. 2g, Extended Data Fig. 4). Despite this, the precise spatial firing pattern of the same cell was uncorrelated between VR and RW (Extended Data Fig. 4).

Intriguingly, certain spiking characteristics were preserved in VR. In RW, neurons generated long spike-sequences lasting about two seconds as rats traversed through well-defined place fields (Fig. 3a, Extended Data Fig. 1). Surprisingly, even without clearly defined place fields, neurons in VR also fired similarly long spike-sequences, appearing as streaks of spikes (Fig. 3b, Extended Data Fig. 1). We term these long spike-sequences “hippocampal motifs,” which we identified as time periods in which a neuron achieved a peak firing rate of at least 5Hz, and maintained a firing rate above 10% of that peak for 300ms. All individual motifs from a cell were aligned around their center of mass and aggregated together to obtain the cell’s motif-field (Fig. 3c, d, see Methods).

**Figure 3.**
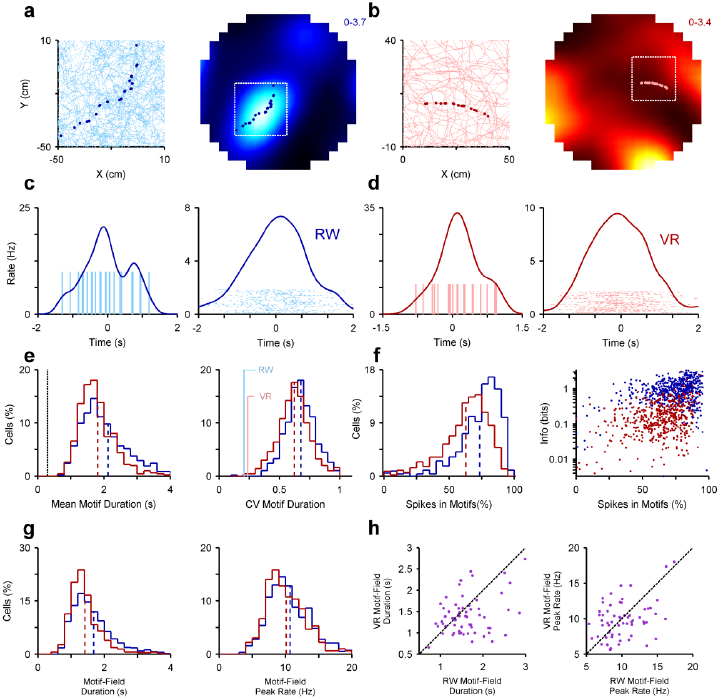
Similar hippocampal motifs and motif-fields in VR and RW. **a)** Spike positions of an example motif from a cell overlaid on a segment of the rat’s trajectory (left) and firing ratemap (right) in RW. **b)** Similar plot as A, but in VR. **c)** Left: Motif firing rate as a function of time and individual spike times (vertical lines) for the same motif as in A. Right: Motif-field firing rate as a function of time. Spikes from individual motifs are depicted in the raster plot, aligned around motifs’ centers of mass to form the motif-field. **d)** Same as C but in VR. **e)** Left: The mean motif durations within each cell were comparable in RW (2.12±0.04s, n=578) and VR (1.81±0.03 s, n=493) but slightly smaller (15%, p<10 ^−10^). The shortest allowed motif duration (dotted vertical black line) was much smaller than the ensemble average. Right: The coefficients of variation (CV) of motif durations within each cell were comparable in RW (0.68±0.01) and VR (0.62±0.01), but slightly lower in VR (8%, p<10 ^−10^). Both were much greater than the CV of the distributions in the left panel (solid vertical lines). **f)** Left: While a majority of spikes were contained within motifs in RW (73.54±0.02%) and VR (62.80±0.03%), there was a significant reduction in VR (15%, p<10 ^−10^). Right: In both RW and VR, the percentage of spikes in motifs was significantly correlated with spatial information content of a neuron (RW r=0.26, p<10^−5^; VR r=0.28, p<10^−5^). **g)** Left: Motif-field durations in VR (1.42±0.02s) were similar but slightly reduced (16%, p<0.05) compared to RW (1.68±0.04s). Right: Peak firing rates of motif-fields in VR (10.13±0.14Hz) were slightly smaller (p<0.05) than in RW (10.69±0.14Hz). **h)** Left: For cells active in both environments on the same day, motif-field duration was correlated between RW and VR (r=0.30, p<0.05). Right: Motif-field peak firing rate had a similar correlation (r=0.42, p<10^−3^).

Motif properties were comparable in the two worlds (Extended Data Fig. 5). The mean motif durations were comparable in RW and VR (Fig. 3e) and motif durations were much longer than expected by chance (Extended Data Fig. 3). Across all cells, mean motif durations displayed small variability (Fig. 3e), which suggests that dorsal CA1 neurons share the same motif time scale on the population level. For any given cell the motif durations were quite variable in RW which could be due to a varying amount of time spent within the place field in each traversal. Curiously, the motif durations were equally variable in VR (Fig. 3e) even though there was little spatial selectivity. Further, in both worlds a majority of spikes were contained within these motifs (Fig. 3f), far greater than expected by chance (Extended Data Fig. 3). Neurons with a larger fraction of spikes within motifs had greater spatial selectivity (Fig. 3f) and mean firing rates (Extended Data Fig. 5). This demonstrates a complex relationship between these three measures, since information content was in fact inversely correlated with firing rate (Fig. 2e). Spiking within motifs, as opposed to isolated spiking, may therefore serve to group otherwise random and non-informative spikes into more informative clusters.

Analysis of motif-fields (Fig. 3c, d) showed similar results, with motif-fields having similar durations in VR and RW (Fig. 3g). The peak rates within motif-fields were also comparable in the two worlds (Fig. 3g), in contrast to the smaller peak rates in spatial ratemaps in VR (Fig. 2a). Neurons active in RW and VR in the same day had motif-fields with similar durations and peak firing rates in the two worlds (Fig. 3h).

We then tested if the motifs showed a temporal code. Hippocampal neurons’ spikes within place fields show theta phase precession—where spikes occur at earlier phases of theta rhythm on subsequent theta cycles—on one-dimensional tracks in RW^4–6^ and VR^7–9^, and in two-dimensional RW environments^10^. Due to the absence of clear place fields in VR we quantified the quality of phase precession within motif-fields by computing the circular linear correlation (see Methods) between the time spent within the motif-field and theta phase of spikes^22^. In the RW 87% of neurons showed significant phase precession within motif-fields. This number was reduced to 45% in VR, but was still far greater than expected by chance (Extended Data Fig. 3, see Methods). For cells with significant precession, the quality of phase precession was comparable in both worlds, although slightly reduced in VR (Fig. 4b). To further test theta scale temporal coding, and to include the data from all cells, we computed the difference between the period of theta modulation of spikes and the LFP theta period ^4^^,^^10^^,^^23^. In both RW and VR this difference was significantly greater than zero (Fig. 4c). Further, a majority of cells in RW (95%) and VR (80%) had longer LFP theta period than spike theta period indicative of intact temporal coding in VR. This is especially notable because the structure of LFP theta was significantly different between the two worlds, with greater peak theta power and reduced theta frequency in VR (Extended Data Fig. 6). The preferred theta phase of neurons was significantly different and more variable in VR than RW (Fig. 4d). Despite these differences, neurons showed identical degrees of theta phase locking in both worlds (Fig. 4e).

**Figure 4.**
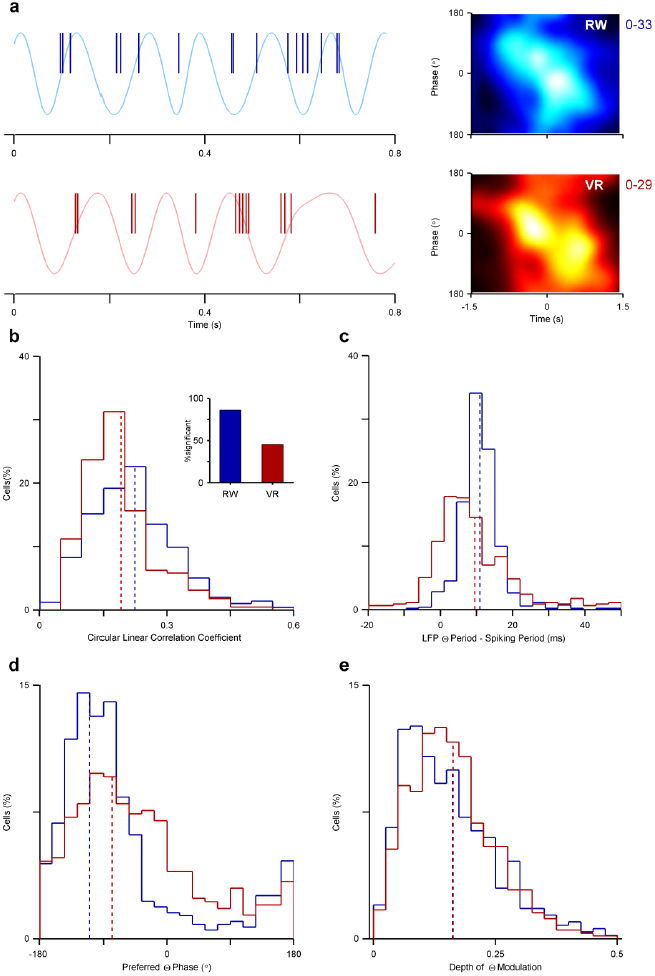
Intact but variable phase coding in VR. **a)** Left: Sample LFP theta traces in RW (top) and VR (bottom) recorded from the same electrode on the same day. Spikes from the same cell (vertical lines) in RW and VR occur at earlier phases on subsequent theta cycles. Right: Motif-fields in RW and VR show clear phase precession. **b)** 87% and 45% of the cells showed significant phase precession in RW and VR respectively (as shown in bar graph). For these, the quality of phase precession in VR cells (0.20±0.01, n=227) was reduced (37%, p<10^−5^) compared to RW (0.25±0.01, n=516). **c)** Difference in LFP theta period and spiking theta period, computed from the autocorrelation of LFP and of spikes shows comparable, although more variable temporal coding in VR (9.49±0.54ms) compared to RW (10.88±0.24ms). **d)** The preferred theta phase of spikes was shifted closer to theta peak (28%, p<10^−3^) in VR (-77.92±0.01°) and more variable in VR (SD=64.04°) compared to RW (-108.29±0.00°, SD=51.25). **e)** The degree of phase locking (depth of modulation) was identical (P>0.05) in VR (0.16±0.11) and RW (0.16±0.09).

These results provide the first measurements of hippocampal CA1 neuronal activity in a two-dimensional environment where only DVC provide reliable spatial information. We found three key results: a profound loss of spatial selectivity in VR; comparable motif dynamics in VR and RW; and comparable temporal coding within motif fields in both worlds. We hypothesize that intrinsic mechanisms within the entorhinal-hippocampal network generate temporally coded motifs that require coherent multisensory inputs for spatial localization as follows.

The large loss of spatial selectivity, stability, and neural activity in VR suggests that DVC alone are insufficient to generate spatial selectivity, which is in contrast to the common finding that DVC govern the spatial localization of place fields^2^^,^^3^. Inputs from multiple sensory modalities are coherent in RW^12^^,^^13^ but not in VR, which could allow the rapid formation of associations via Hebbian synaptic plasticity, thus establishing a stable spatial representation^24^. This can explain why spatial selectivity was intact when the rats ran on one-dimensional VR^7–9^ since self-motion and internal cues were always coherent with DVC, which is not the case in two dimensional random foraging in VR. Even in a two-dimensional RW, a change in coherence between self-motion cues and DVC results in a change in spatial activity pattern and directionality^25^. This is also consistent with the observation that maze-cleaning between sessions induced place field instability and remapping^12^. The results are unlikely to arise solely due to diminished vestibular inputs in VR because vestibular lesions caused significant behavioral deficits, reduction in theta power and unaltered peak firing rates^15^^,^^26^, all of which are in contrast to our data.

Despite the loss of spatial selectivity in VR, neurons had a comparable tendency to fire long spike-sequences or motifs, similar to those in RW, supporting an intrinsic, network mechanism of motif generation. This is further supported by the small variability in motif durations on a population level compared to a neuronal level, together with correlated motif-field properties between worlds. Additionally, the fraction of spikes contained within motifs was comparable in RW and VR despite reduced spatial selectivity.

Intact theta scale dynamics in VR motif fields suggests that intrinsic, network mechanisms also govern temporal coding. Shifted preferred theta phase and its increased variability could arise via a rate-phase transformation and reduced excitatory drive in VR due to lack of coherent multisensory inputs. The intact theta scale dynamics without spatial selectivity in VR shows that the hippocampal temporal code can exist without the place code.

Unlike sensory cortices where neurons generate brief responses, motif activity could be an intrinsic property of the entorhinal-hippocampal network which could explain their emergence in diverse ways including hippocampal place cells^7–9^, entorhinal cortical grid cells^23^, episode or time cells during wheel or treadmill running^16^^,^^18^, and neural activity during free recall^27^ and REM sleep^28^.

Motifs could originate from several parts of the entorhinal-hippocampal network. The recurrent CA3 network offers a potential candidate. Alternatively, the motifs could arise in the medial entorhinal cortex where neurons show motif-like activity lasting a few seconds and robustly driving CA1, even in anesthetized or sleeping animals^29^. Consistently, sustained spiking in consecutive theta cycles was reduced, indicative of diminished motifs, in a transgenic mouse with diminished distal dendritic inputs, which typically originate in the entorhinal cortex^30^. The motif-field durations along the dorso-ventral axis of the entorhinal-hippocampal circuit could also be influenced by the temporal integration properties of the h-current along that axis^31^. We propose that behaviorally relevant patterns of activity may thus be constructed as internally generated, temporally coded and cue-localized motifs from spikes occurring at much shorter time scales. This could function to help bridge the gap between the rapid dynamics of neural and synaptic activations and the longer time scales of behavior^6^.

## Methods Summary

Three adult male Long-Evans rats were trained to forage for randomly scattered rewards in two-dimensional RW and VR environments. The environments had identical dimensions (200cm diameter circular platform at the center of a 300×300cm room) and DVC. Electrophysiological data from dorsal CA1 were obtained using hyperdrives with 22 independently adjustable tetrodes^9^. Spike extraction and sorting were done offline using custom software. A detailed description of the methods and analysis is available in the Methods section.

## Acknowledgements

We thank N. Agarwal for help with electrophysiology, B. Popeney for help with behavioral training, F. Quezada for help with behavioral training and spike sorting, B. Willers for help with the analyses, P. Ravassard and A. Kees for help with surgeries, and D. Aharoni for help with hardware. This work was supported by grants to MRM from: NIH 5R01MH092925-02 and the W. M. Keck foundation.

## Online Methods

Materials and methods were similar to those described recently ^9^^,^^21^.

### Subjects

Data were collected from three adult male Long-Evans rats (approximately 3.5 months old at the start of training), individually housed on a 12 hour light/dark cycle and food restricted (15-20 g of food per day) to maintain body weight. They were allowed an unrestricted number of sugared water rewards in VR but a restricted amount of water (30-40ml of water per day) after the behavioral session to maintain motivation. All experimental procedures were approved by the UCLA Chancellor’s Animal Research Committee and were conducted in accordance with USA federal guidelines.

### Random Foraging in RW and VR

The experimental room, the VR apparatus, and basic behavioral training were identical to those described recently^9^^,^^21^. In RW, a 200cm diameter and 50cm high platform was placed at the center of a 300×300cm room with distinct visual cues on the four walls (Fig. 1a). Rats were trained to forage for randomly scattered food rewards on the platform. The VR room had identical size and DVC, and rats foraged for randomly located rewards on a platform of the same size as in the RW room. Rewards in VR were in the form of sugar water dispensed through reward tubes placed directly in front of the rats. The reward locations were hidden and 60cm in diameter. Entry into the reward locations triggered the appearance of a white dot of the same size on the platform in addition to a reward tone and sugar water delivery. At each reward location rats could receive a maximum of five sugar water rewards. Motion parallax between the virtual elevated table and the floor underneath indicated the virtual edge of the platform. Movement beyond the platform edge resulted in no change in visual scene. Rats quickly learned to avoid or turn away from the virtual edges (Fig. 1a). It took about three weeks of handling and pre-training and two weeks of VR training for rats to do the random foraging task efficiently. Rats were trained on the RW task after implantation. Two rats were run in both RW and VR every day. To verify that exposure to both worlds on the same day was not playing a role in neural responses, another rat never ran in both RW and VR on the same day. Further, the order of running on VR and RW on the same days was randomized. No qualitative differences were found between these conditions and hence all data were combined.

### Surgery, Electrophysiology and Spike Sorting

These procedures were identical to those described earlier^9^. Briefly, once the rats reached performance criterion they were anesthetized using isoflurane. Custom-made hyperdrives containing up to 22 independently adjustable tetrodes that targeted both left and right dorsal CA1 were implanted. Rats were allowed to recover from surgery for one week after which the tetrodes were gradually advanced to area CA1, detected online by clear presence of sharpwave ripple complexes. Spike and LFP data were recorded at 40Khz. Spikes were extracted and sorted into individual units using custom software. Classification of single unit cell type was performed using the same methods as previously described^9^. When rats ran in both VR and RW on the same day, the same cells were identified by overlaying cluster boundaries from both sessions, and identifying clear overlaps. If cell identities were unclear due to electrode drift the data were discarded from the same cell analysis.

### Statistics

Offline analyses were performed using custom MATLAB codes. Tests of significance between linear variables and circular variables were done using the nonparametric Wilcoxon rank-sum test and Kuiper test respectively. Tests of significance for the mean values of distributions being different from zero were performed using the nonparametric Wilcoxon signed-rank test. To compute circular statistics, CircStat toolbox was used^22^. All ensemble averages are in the form mean ± SEM unless otherwise stated.

### Quantification of Ratemaps

Theta rhythm is interrupted and behavior is uncontrolled when rats paused to consume rewards or to groom. Hence these periods were excluded and only data recorded during periods of active locomotion (running speed > 5cm/s) were used. A cell was considered active if its mean firing rate exceeded 0.2Hz during locomotion and was thus included in the analysis. Spatial firing rates were computed using occupancy and spike histograms with 10×10cm bins smoothed with a 15cm two-dimensional Gaussian smoothing kernel. Bins with very low occupancy relative to the experimental session were excluded to avoid artificially high firing rates. The spatial information content, sparsity and coherence of the ratemaps were computed using methods described recently^9^. To determine the stability of ratemaps, firing rates were computed in the first and second halves of the session separately. The bin-by-bin correlation between the ratemaps in the two halves provided a measure of ratemap stability. To obtain the similarity of ratemaps of the same cell in RW and VR we computed the correlation of firing rates and computed statistical significance by comparing it against correlations when cell identities were shuffled.

### Detection of Motifs

To detect motifs a method similar to the one used for detecting place fields on a one dimensional track was used. We constructed a spike train, a vector of data whose length was equal to the period of experimental session by binning the spikes for which the running speed was greater than 5cm/s. This spike train was smoothed using a 200ms Gaussian smoothing kernel and transformed to firing rate by dividing by the bin duration. Peaks where firing rate exceeded 5Hz were detected and marked as candidate motifs. The boundaries of a motif were defined as the points where the firing rate first dropped below 10% of the peak rate (within the motif) for at least 250ms (two theta cycles). If the time-lag between the first and last spike in the putative motif, called the duration of the motif, exceeded 300ms, this sequence was considered a valid motif and was included in the analysis.

### Construction of Motif-fields

The center of mass within each motif was computed using the firing rate to determine the center of the motif. This value was subtracted from the spike times within the motif to center them around zero. This procedure was repeated for all motifs and the centered motifs were aligned to obtain a motif-field for a neuron. The firing rate as a function of time within motif-field was calculated as the number of spikes within each temporal bin divided by the total amount of time in that bin, smoothed by a 200ms Gaussian smoothing kernel. Motif-field duration was defined as twice the weighted standard deviation of the motif firing rate, i.e. the width of the distribution.

### Theta Period and Phase Precession

Similar to the methods described previously^9^, each LFP was filtered between 4 and 12 Hz using a 4^th^ order Butterworth filter. Theta period was computed by detecting the peak between 50 and 200ms in the filtered LFP autocorrelation for epochs when the running speed was above 5cm/s. Spiking theta period was calculated by computing the spike train autocorrelation, smoothing it with a 15ms wide Gaussian smoothing kernel, and detecting the peak. Quality of phase precession within a motif-field was defined as the circular linear correlation coefficient (CLCC)^9^ between spike phases and latency of spike timing with respect to the motif center.

### Control Analysis for Motifs

To estimate which motif properties can arise purely by chance, surrogate motifs for each neuron were generated as follows. The mean firing rate during locomotion and the depth of theta modulation were computed for each neuron. Surrogate activity was generated using a Poisson distributed and theta modulated spike train with the same mean firing rate and depth of theta modulation as the experimentally measured neuron. Motifs, motif-fields, and their properties were computed using procedures described above. This procedure was repeated 50 times for each neuron to generate a null distribution.

### Control Analysis for Precession

To determine how much phase precession can occur by chance, for each cell its entire spike time series was shifted by a random amount of time between 10 and 20 seconds, repeated 50 times. This conserves the temporal structure of the spikes but, due to variations in theta frequency, randomizes spike phases. Motif-fields were then detected and quality of phase precession was quantified as described earlier.

---

**Extended Data Figure 1.**
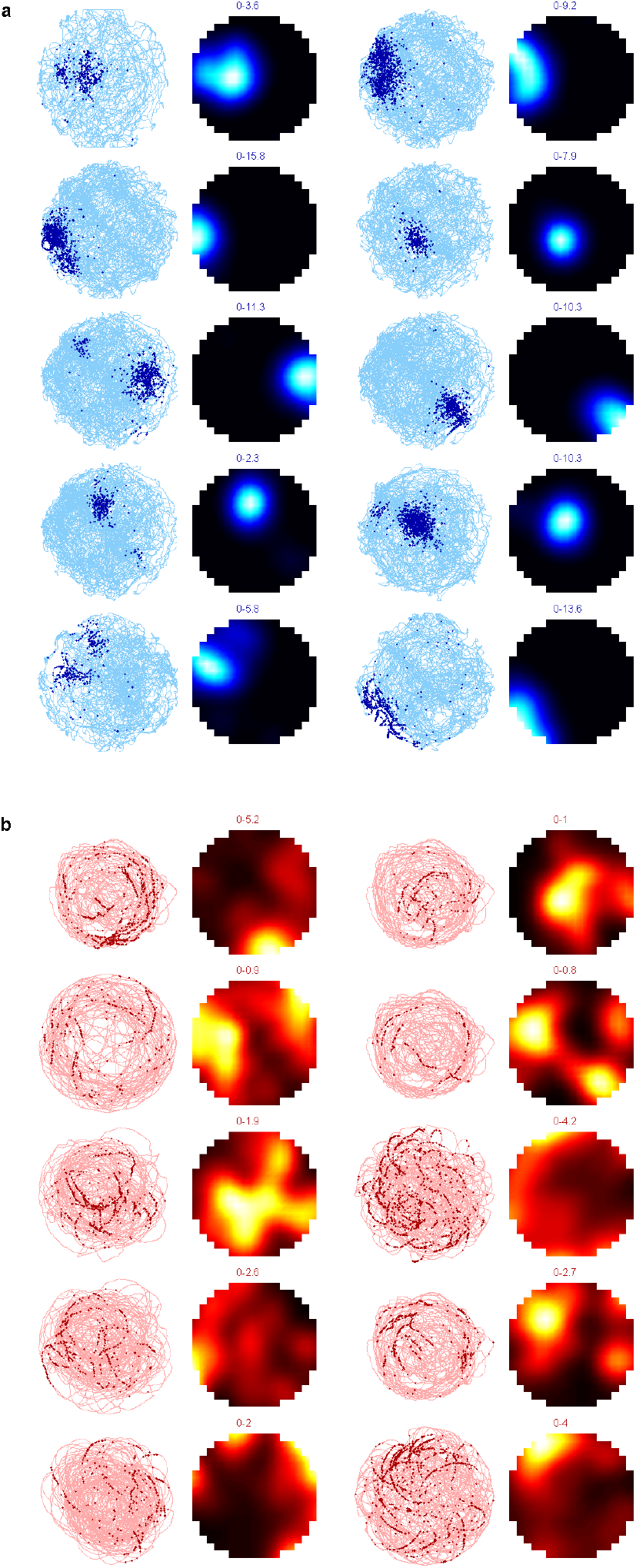
Additional examples of place cells in VR and R. **a)** Rat trajectory and spike positions for different neurons and corresponding firing ratemaps in RW. **b)** Same as A but in VR, showing long streaks of spikes, or putative motifs. Numbers indicate firing rate range.

**Extended Data Figure 2.**
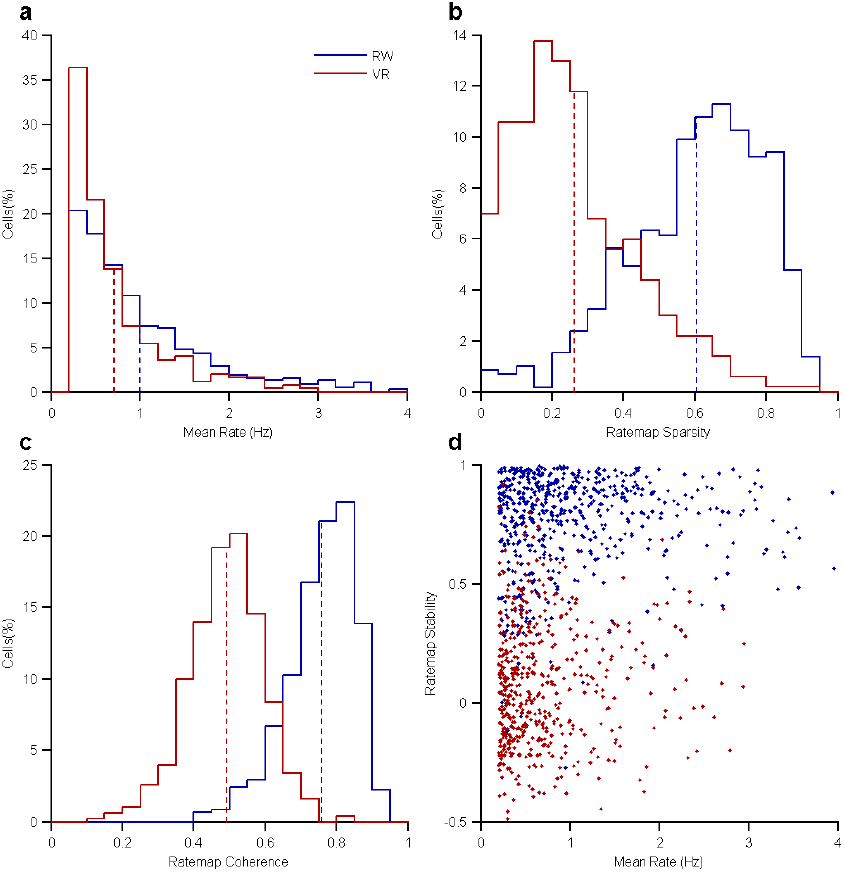
Reduced mean firing rates, ratemap sparsity, coherence in VR. **a)** Mean firing rates were 29% (p<10^−10^) lower in VR (0.71±0.02Hz) than in RW (0.99±0.03Hz). **b)** Ratemap sparsity, a measure of spatial selectivity, was also greatly (57%, p<10^−10^) reduced in VR (0.26±0.01) compared to RW (0.60±0.01). **c)** Ratemap coherence was 35% (p<10^−10^) reduced in VR (0.49±0.01) compared to RW (0.76±0.01). **d)** At all mean rates, stability was lower in VR compared to RW. Stability was not correlated with mean firing rate in either world (r=0.05, p>0.05 and r=0.08 and p>0.05 in RW and VR respectively).

**Extended Data Figure 3.**
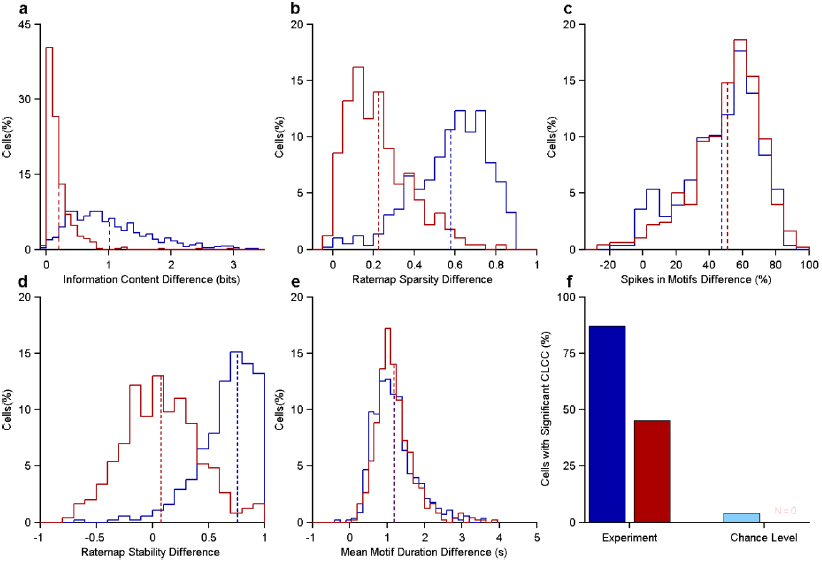
Comparison of spatio-temporal properties of experimental data with chance level. Surrogate spike trains with similar mean firing rates and theta modulation index as experimental data (see Methods) were generated for each cell in VR and RW. These were repeated 50 times for each cell and the mean value of the spatio-temporal property measure computed by averaging the numbers of these 50 repetitions. These mean values were subtracted from the numbers obtained for the experimental data which yielded the difference between the experimental values and the values expected by chance. The values obtained by chance were so small that the difference distribution for all measures was nearly identical to the actual experimental measures indicating that all were highly significant. In the control analysis for phase precession (see Methods), the number of significant CLCCs were computed. **a)** Spatial information content was significantly greater than chance level in VR (0.20±0.01 bits, p<10^−10^) but the difference was greater in RW (1.02±0.03 bits, p<10^−10^), and the two distributions were significantly different (p<10^−10^). **b)** Similar to information content, ratemap sparsity was significantly greater than chance level in VR (0.22±0.01, p<10^−10^) but the difference was greater in RW (0.58±0.01, p<10^−10^). These two distributions were significantly different (p<10^−10^). **c)** The occurrence of motifs was significantly above chance in VR (0.51±0.01, p<10^−10^) and comparable to RW (0.48±0.01, p<10^−10^) and the two distribution were similar (p>0.05). **d)** The stability was close to chance level in VR (0.08±0.01, p<10^−10^) but significantly above chance in RW (0.76±0.012, p<10^−10^). **e)** The mean motif durations were significantly longer than the motifs generated from surrogate spike trains in both worlds (0.50±0.03 s, p<10^−10^ and 0.61±0.02 s, p<10^−10^ in RW and VR respectively). **f)** In the time-shifted spike trains, there were only four significant phase precession CLCCs in RW and none in VR.

**Extended Data Figure 4.**
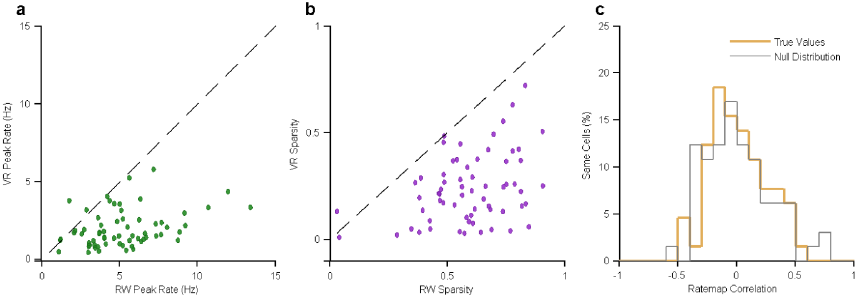
Comparison of activities of cells active in both VR and RW on the same day. **a)** The peak firing rate of the same cell in RW and VR was reduced but significantly correlated (r=0.30, p<0.05). **b)** Sparsity of the same cell in RW and VR was also reduced but correlated (r=0.37, p<0.01). **c)** Despite positive correlations in peak rate, information content (Fig. 2G) and sparsity, the ratemaps of the same cells in RW and VR were uncorrelated (true values, p>0.1) and not different from the correlations obtained by shuffling the cell identities (null distribution, p>0.5).

**Extended Data Figure 5.**
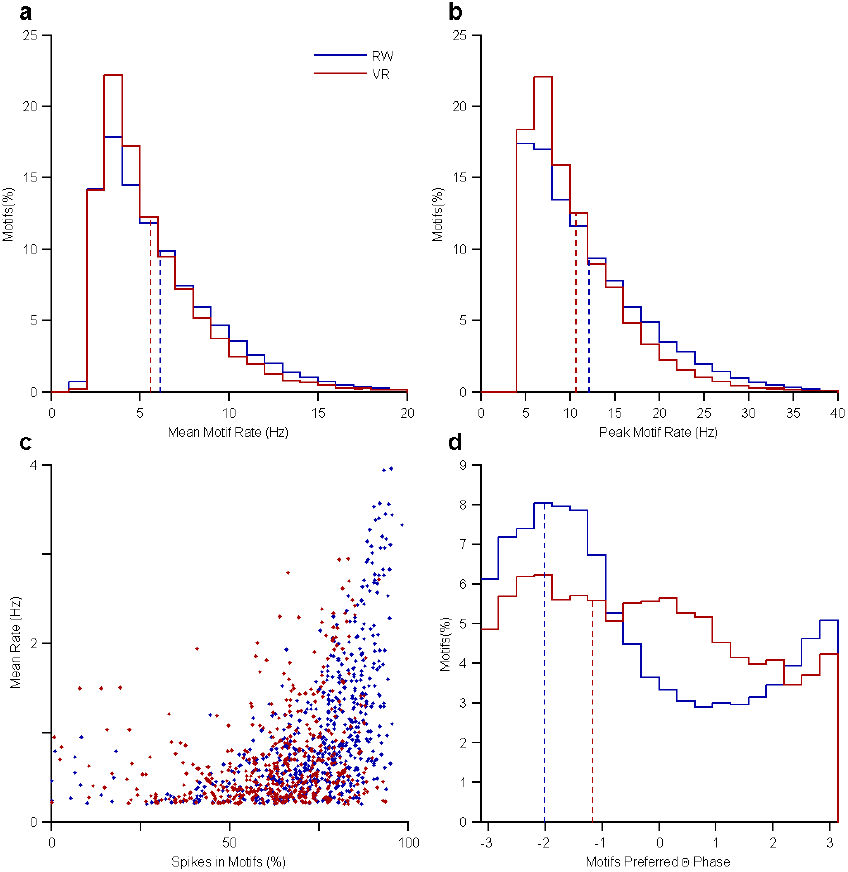
Spatio-temporal properties of individual motifs in VR and RW. **a)** Mean firing rates within motifs in VR (5.61±0.02Hz) were slightly smaller (8%, p<10^−10^) than in RW (6.13±0.02Hz). **b)** Similar reduction (13%, p<10^−5^) was observed in peak rate within motifs in VR (10.68±0.04Hz) compared to RW (12.22±0.06Hz). **c)** There is significant correlation between mean rate and the percentage of spikes that occurred within motifs in RW (r=0.52, p<10^−10^) and VR (r=0.29, p<10^−10^). **d)** Preferred theta phase within the motifs in VR (-67.26±76.95°) was shifted towards theta peak and more variable (p<10^−3^) compared to RW (-115.62±69.84°).

**Extended Data Figure 6.**
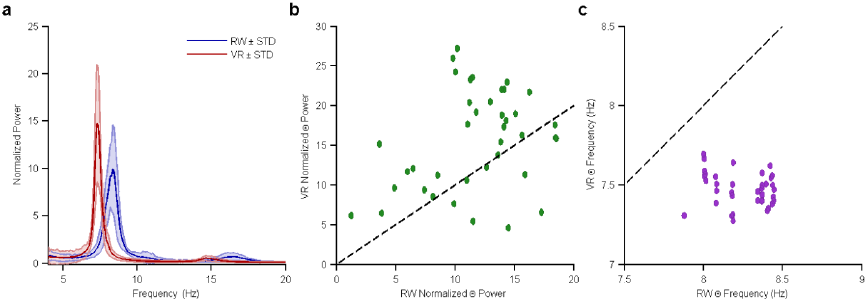
Increased Theta Power but Reduced Theta Frequency in VR. To further examine the dynamics of LFP theta, we investigated the LFPs recorded from the same electrode on the same day in both worlds without any electrode movement in between the two sessions. Analysis was further restricted only to data when rats ran at speeds greater than 5cm/s to eliminate contamination by variable periods of stopping when theta is reduced. LFP power spectrum was first computed and then restricted to the range of 4-20 Hz. In order to compare data from different sessions, power spectrum from each electrode was normalized by the mean power on that electrode in VR and RW over the same range. **a)** Normalized power between 4-20 Hz, averaged over all the LFP (n=39) in RW and VR shows a clear shift in theta power and frequency between the two environments. **b)** Peak theta power is significantly increased in VR (p<10^−5^). **c)** Theta frequency is significantly lower in VR (p<10^−5^).

